# Niche cytoskeletal architecture is required for proper stem cell signaling and oriented division in the *Drosophila* testis

**DOI:** 10.1101/2024.06.04.597489

**Authors:** Gabriela S. Vida, Elizabeth Botto, Stephen DiNardo

**Author notes:** Corresponding Author/ Lead contact Correspondence. Morsani College of Medicine, University of Southern Florida, 560 Channelside Dr. Tampa, FL 33602.

## Abstract

Stem cells are critical to repair and regenerate tissues, and often reside in a niche that controls their behavior. The *Drosophila* testis niche has been a paradigm for niche-stem cell interactions and is used here to address the cell biological features that maintain niche structure and function during its steady-state operation. We report enrichment of the Myosin II (MyoII) and a key regulator of acto-myosin contractility (AMC), Rho Kinase (ROK), within the niche cell cortex at the interface with germline stem cells (GSCs). Compromising MyoII and ROK disrupts niche architecture, suggesting that AMC in niche cells is important to maintain the reproducible structure of this niche. Furthermore, defects in niche architecture cause changes in stem cell function. Our data suggest that the niche signals less robustly to adjacent germ cells, yet the disrupted structure permits increased numbers of cells to respond to the signal. Finally, compromising Myo II in niche cells leads to an increase in mis-oriented centrosomes in GSCs as well as defects in the centrosome orientation checkpoint. Ultimately, this work identifies a critical role for AMC-dependent maintenance of niche structure to ensure a proper complement of stem cells with correct execution of stem cell divisions.

**Summary statement:** Actomyosin contractility regulated niche architecture is critical for proper signaling and oriented division of germline stem cells.

## Introduction

Stem cells are critical in supporting and repairing the tissues in which they reside, and are often maintained by a niche, a specialized microenvironment. The niche is important for sending cues that allow stem cells to maintain their identity and regulate the balance between self-renewing and differentiating outcomes. Imbalance of these cues can lead to tumor formation or tissue degeneration (Ohlstein et al., 2004; Morrison and Kimble, 2006; Morrison and Spradling, 2008). Therefore, the proper function of the niche is required for tissue homeostasis.

Stem cell niches across organisms and tissues often adopt a reproducible shape, structure, or organization. For example, the mammalian intestinal epithelium organizes its niche into the base of crypts. This affords a protected environment within which stem cells are maintained, while newly differentiating cells travel out of the crypt and replace cells during intestinal lining turnover (Barker et al., 2007; Sumigray et al., 2018). However, what remains less understood is whether that precise crypt organization is critical for stem cell function. Recent work in intestinal organoids revealed a requirement for crypt invagination and its precise diameter to promote stem cell function (Pentinmikko et al., 2022). While it is speculated that specific niche architectures are critical to regulate signaling and stem cell behaviors, the mechanisms controlling niche structure and the effects of disruption on stem cell dynamics remain unknown.

The *Drosophila* testis is an ideal system to study the potential role of niche structure on stem cell function. The identity of the stem cells and niche cells are known, and their respective behaviors can be directly visualized within the tissue through live imaging. The testis niche is composed of a compact group of quiescent somatic cells that reside at the apex of the testis tube (Hardy et al., 1979). The niche signals to two stem cell populations, the germline stem cells (GSCs) and the cyst stem cells (CySCs)(Kiger et al., 2001; Tulina and Matunis, 2001; Kawase, 2004; Leatherman and DiNardo, 2008; Leatherman and DiNardo, 2010; De Cuevas and Matunis, 2011; Amoyel et al., 2013; Inaba et al., 2015b). Signals from the niche exhibit spatially restricted effects, with pathway activation reaching only the first tier of cells adjacent to the niche and thus maintaining a restricted stem cell pool. For example, niche secretion of the ligand Unpaired (Upd) activates the Jak/STAT pathway only in nearby cells (Kiger et al., 2001; Tulina and Matunis, 2001). This contained signaling is critical for stem cell maintenance and attachment at the niche and thus allows differentiation of germ cells once displaced from the niche.

In many tissues, the placement of the niche imparts an overall directionality to differentiating cells as they move away from the niche. This is the case in the *Drosophila* testis because the stem cell divisions are invariably oriented such that differentiating daughters are displaced away from the niche (Yamashita, 2003; Yamashita et al., 2007). The mechanism controlling this is most understood for GSCs where an intrinsic regulatory network controlling division plane has been identified. Multiple factors, including E-cadherin, Bazooka/Par-3 and APC, are enriched in the GSC cortex adjacent to niche cells, where they anchor one of the centrosomes (Yamashita, 2003; Inaba et al., 2010; Inaba et al., 2015a). Centrosome anchoring orients the eventual mitotic spindle so that division occurs orthogonally, generating one daughter cell remaining attached to the niche and one daughter pushed away. Such orthogonal divisions seem critical in GSCs since a centrosome orientation checkpoint (COC) exists, and is employed specifically in the GSCs to monitor centrosome anchoring (Venkei and Yamashita, 2015). If centrosomes are misoriented, the COC arrests the GSC cell cycle in G2 until centrosomes are properly positioned (Cheng et al., 2008; Venkei and Yamashita, 2015). Currently, only GSC-intrinsic factors regulating the COC have been identified, including Centrosomin and Par-1 (Inaba et al., 2010; Yuan et al., 2012a). However, the precise mechanisms controlling centrosome anchoring and how the GSC cortex adjacent to the niche cell must be specialized to sense proper anchoring is not well understood.

While little is known about how niche architecture is maintained in the adult testis, substantial work has been done to understand how the niche adopts this architecture during its initial formation. The structure of the steady-state niche is initially formed in the embryo where a subset of somatic gonadal cells are specified to become niche cells, migrate and assemble loosely at the anterior of the gonad, and then undergo compaction to achieve a spherical shape (Le Bras and Van Doren, 2006; Anllo et al., 2019; Anllo and DiNardo, 2022; Warder et al., 2023). Work from our lab has shown that actomyosin contractility (AMC) is required during the compaction process to shape the niche. Inhibition of AMC or the AMC regulator Rho Kinase (ROK) in the gonad disrupts compaction and ultimately generates defective niche architecture. In parallel work analyzing snRNA-seq data from the steady-state testis, we found that ROK was predicted to be enriched in the niche compared to other cells in the testis (Li et al., 2022; Raz et al., 2023). This finding in conjunction with our lab’s work on gonadal niche assembly, suggested a role for AMC beyond formation to include maintenance of niche structure.

Previous analyses disrupting niche structure also altered niche cell and/or fate. For example, knockdown of retinoblastoma (Rb) in the niche allowed for huge changes in niche size and structure, but also resulted in niche cells exiting quiescence and growing in number (Greenspan and Matunis, 2018). Additionally, ablation of CySCs can induce niche cells to exit quiescence and can result in ectopic niche formation after recovery from CySC ablation (Hétié et al., 2014). Exiting the normally quiescent state, changes to niche cell number, and/or alterations in fate, all make it difficult to assess how niche architecture might contribute to niche function in governing stem cell behavior.

Here we describe a requirement for AMC in maintaining niche structure in the adult testis niche. We show that reduction in AMC causes changes to the polarized localization of adhesion proteins between niche cells, concomitant with progressive disassembly of the niche. These changes in niche structure occur without increased niche cell numbers, loss of niche quiescence or altered niche cell identity. Thus, we could now ask if maintenance of niche structure is necessary for proper stem cell regulation in the absence of confounding factors. Importantly, loss of proper niche structure disrupts restriction of Jak/STAT signaling, resulting in increased numbers of STAT+ germ cells, most with decreased activity of the pathway. Lastly, we have identified the first GSC-extrinsic control of the COC, as we find that disrupted niche structure prevents proper execution of oriented divisions in GSCs. Taken together, we elucidate mechanisms required for long-term maintenance of niche structure and identify the critical importance of niche architecture in proper regulation of stem cell.

## Results

### AMC components are enriched in the adult testis niche

While significant work has been done studying niche formation in the embryo, much less is known about maintenance of niche structure in the adult tissue. To address this, we first examined the niche in three dimensions (3D) to determine if niche architecture remains similar to the spheroid structure established during embryogenesis (Warder et al., 2023). Outlining niche cell membranes using either Fasciclin III (Fig 1A’,1B) or E-Cadherin (Fig 1A’’,1B’), revealed niche structure to indeed approximate a spheroid (Fig 1C, see Materials and Methods), similar to that observed upon niche formation and consistent with the classical description of this niche based on electron microscopy (Hardy et al., 1979).

**Figure 1:**
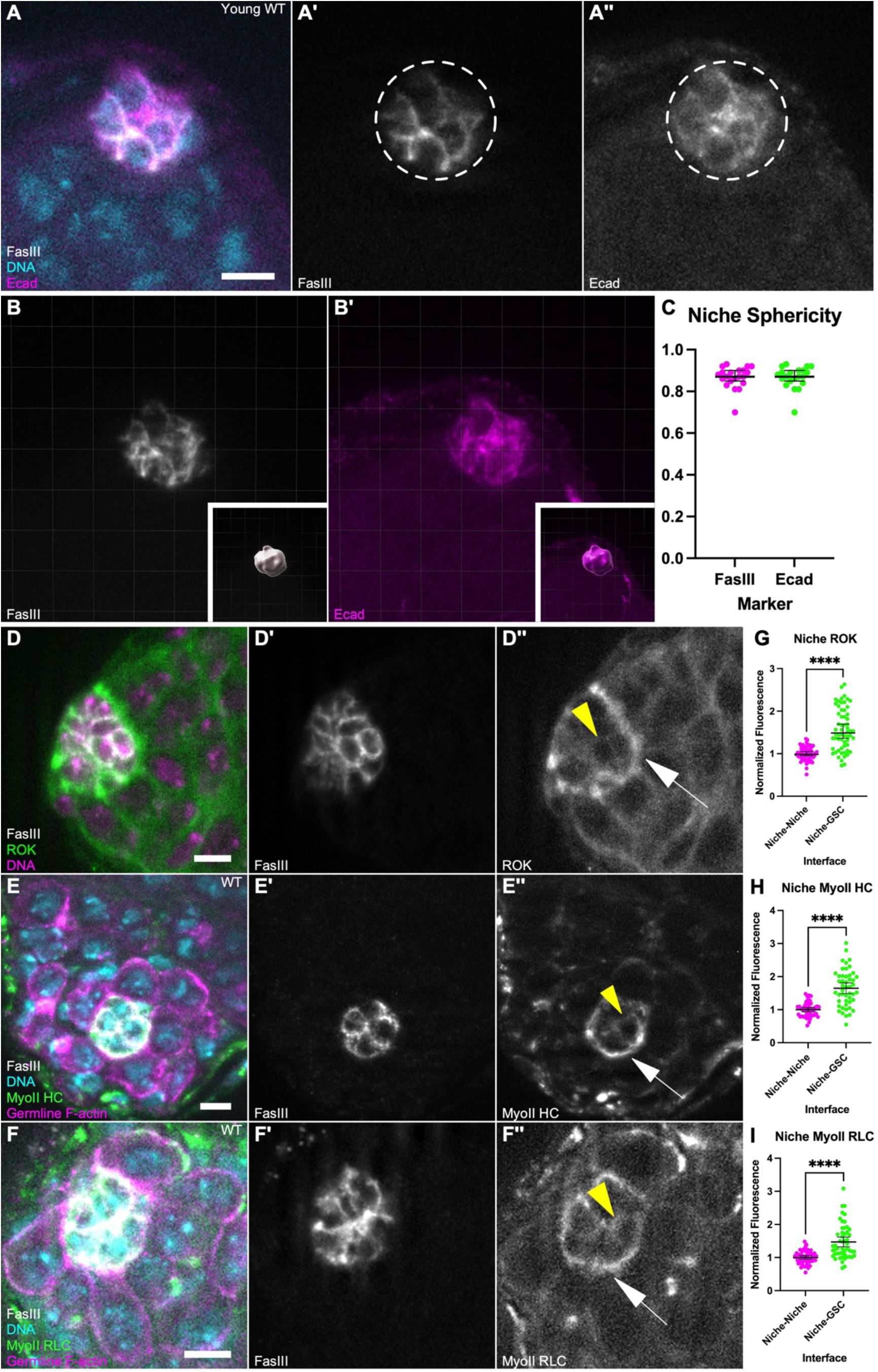
AMC components are enriched in the testis niche. (A-B) Immunostained young w1118 adult testis (A) then displayed in 3D (B,single plane) (A) FasIII (white), E-cadherin (magenta); Hoescht (cyan). (A’,B) FasIII only; (A’’,B’) E-cadherin only; (A) niche represented by white dotted line (B) insets show surface imposed on niche. (C) Quantification of sphericity using FasIII and E-cadherin as markers; sphericity of niche approaches 1. (D-F) Immunostained young testis; (D) ROK::GFP, FasIII (white), Hoescht (magenta); (E) MyoIIHC::GFP, nanos-lifeact::tdTom, FasIII (white), Hoescht (cyan); (F) MyoIIRLC::3xGFP, nanos-lifeacttdTom, FasIII (white), Hoescht (cyan); (D’,E’,F’) FasIII only; (D’’) ROK only; (E’’) MyoIIHC only; (F’’) MyoIIRLC only; (D’’,E’’,F’’) yellow arrowhead points to a niche-niche interface, white arrow points to niche-GSC interface. (G-I) Quantification of enrichment at Niche-GSC interfaces (D’’,E’’,F’’, white arrow) in comparison to Niche-Niche interfaces (D’’,E’’,F’’, yellow arrowhead) for (G) ROK::GFP, (H) MyoIIHC::GFP, (I) MyoIIRLC::GFP (****p<0.0001, Mann Whitney test). Scale bar =5μm.

Our lab recently showed that AMC is critical in achieving this spheroidal shape during niche formation (Warder et al., 2023). To test whether contractility remained important in preserving niche structure, we examined the distribution of a regulator of contractility, ROK, that was predicted to be upregulated in niche cells by snRNAseq data (Raz et al., 2023). A ROK::GFP transgenic reporter exhibited enriched signal at the niche compared to other cells in the testis (Fig 1D-D’’). In addition, by quantifying ROK::GFP fluorescence intensity we found that ROK was polarized to the niche-GSC interface as opposed to the niche-niche interface (Fig 1G). Since ROK governs contractility through its action on Myosin II (MyoII), we next asked whether MyoII was similarly enriched in niche cells. We used fluorescently tagged transgenic lines for either MyoII heavy chain (HC, zipper, Fig 1E-E’’) or MyoII regulatory light chain (RLC, squash, Fig 1F-F’’) and found that both were similarly polarized to the niche-GSC interface (Fig 1H,1I). Therefore, MyoII and a key regulator of contractility are polarized at the niche-GSC interface in the adult testis, precisely located to play a functional role in maintenance of niche structure.

### ROK is required for maintenance of niche shape

To determine whether ROK is important for the preservation of niche shape, we temporally induced expression of an RNA interference (RNAi) against ROK using a niche-specific driver, Upd-GAL4 regulated by Gal80ts. We confirmed specificity and temporal induction of the system by expressing a UAS-GFP under GAL4-restrictive and GAL4-permissive temperatures and observed GFP expression localized to the niche only upon upshift to the restrictive temperature for Gal80ts (Fig S1). For all RNAi treatments, flies were initially grown without knockdown until late pupal stages, allowing normal assembly and function of the niche. Flies were then upshifted to activate GAL4 and initiate knockdown by expression of the target RNAi, along with the UAS-GFP, and assayed after 10 days. After ROK knockdown, niche structure was misshapen in about 45% of niches (Fig 2A-D). A misshapen niche qualifies as a niche that has lost its spheroid structure to adopt more elongate shapes. To quantify the degree of misshaping, we measured niche sphericity, area, and volume in 3D. Niches compromised for ROK exhibited decreased sphericity and increased area and volume in comparison to control niches (Fig 2E,H-I). Importantly, using a nuclear marker, we quantified niche cell number and found no difference between wildtype and ROK knockdown niches (Fig 2F). By contrast, the average internuclear distance among niche cells was significantly increased upon ROK knockdown, suggesting that increased niche area and volume was due to dispersion of niche cells (Fig 2G; see Materials and Methods). Since these niche cells still retained niche-specific Fasciclin III and continued to express Upd-GAL4, these data strongly suggest that, while ROK activity is not necessary for maintenance of niche cell identity, it is essential to maintain proper niche shape.

**Figure 2:**
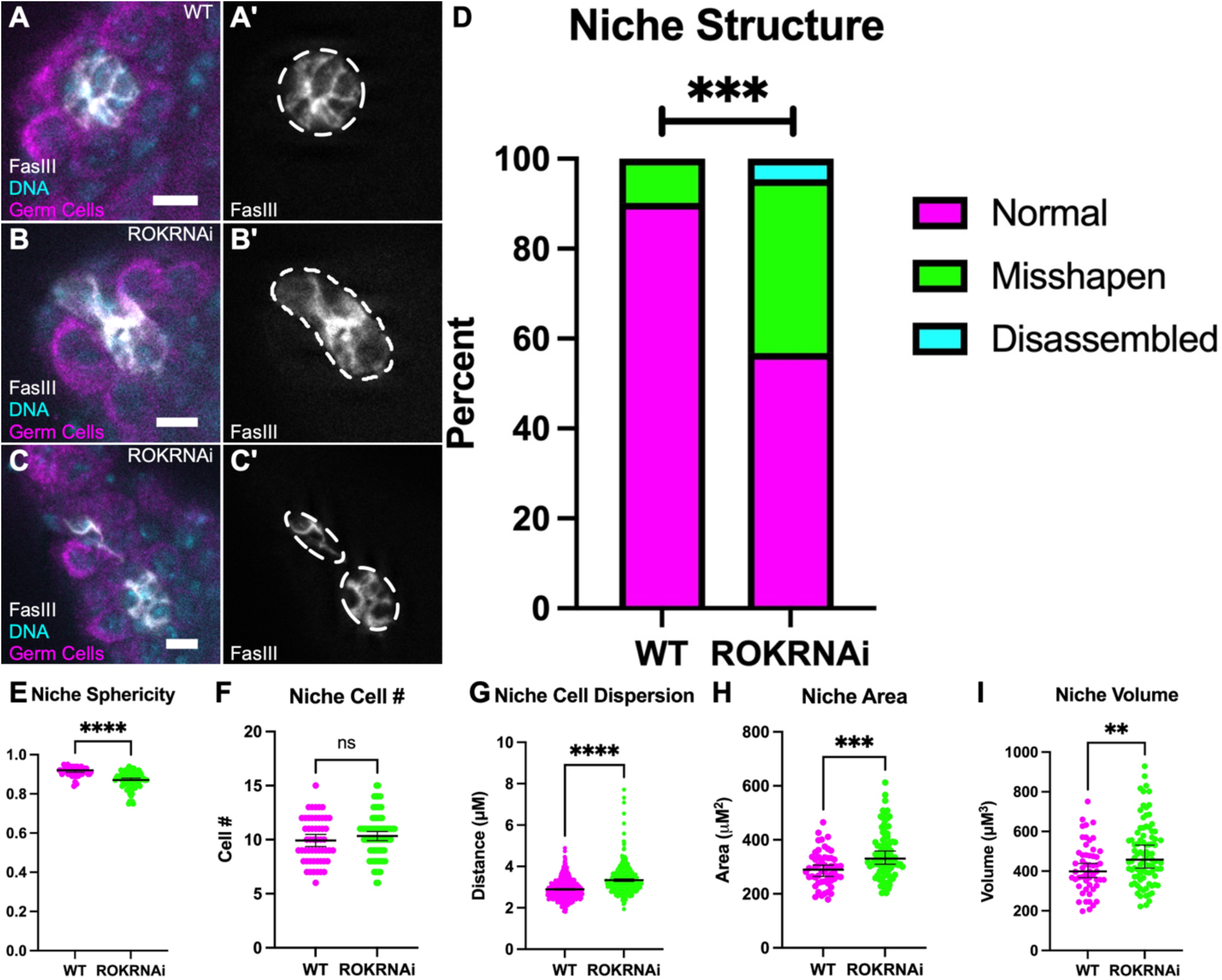
ROK is required for niche shape. (A-C) Immunostained WT testis (A) and ROKRNAi testis where niche is misshapen (B) or disassembled(C); (A,B,C) Vasa (magenta), FasIII (white), Hoescht (cyan); (A’,B’,C’) FasIII only; niche shape represented by white dotted line. (D) Quantification of the niche phenotype shows ROK RNAi niches are severely misshapen and occasionally disassembled into more than one aggregate compared to age-matched control niches (WT: 5/51 niches misshapen, ROKRNAi: 34/88 niches misshapen, 4/88 niches disassembled, ***p=0.0002, Chi-squared test). (E) Quantification of sphericity between age-matched control and ROK RNAi niches (****p<0.0001, Mann-Whitney test). (F) Quantification of niche cell number show no difference between age-matched control and ROK RNAi niches (ns). (G) Quantification of the average distance of a niche cell and its 3 nearest neighbors show increased distance between ROK RNAi niche cells compared to age-matched control (****p<0.0001, Mann-Whitney test). (H-I) Quantification of the niche area (H) and volume (I) shows an increase in ROK RNAi niches compared to WT (***p=0.0001, **p=0.0091, Mann-Whitney test). Scale bar =5μm.

### MyoII is required for niche shape

As a main role for ROK is control of MyoII contractility and given the enrichment of both at niche-GSC interfaces, we next asked whether MyoII is similarly important for preservation of niche shape. We expressed an RNAi against MyoIIHC and quantification of MyoII fluorescence intensity (Zipper) confirmed that protein levels were significantly reduced by RNAi expression in niche cells (Fig 3, compare J” and K”; L). We found that niches depleted of MyoII were severely misshapen compared to controls (Fig 3A-C, Fig S3). Notably, about one third of these niches were split into separate aggregates (Fig 3D). Loss of MyoII consistently led to decreased niche sphericity, and increased area and volume in comparison to wildtype niches (Fig 3E,H-I). Consistent with our results from ROK knockdowns, loss of MyoII in the niche did not alter niche cell number but resulted in greater dispersal of niche cells compared to controls (Fig 3F-G). Importantly, cells comprising the altered niches remained quiescent, as none were positive after a pulse with the S-phase substrate, EdU (Fig S2). These data together reveal a requirement for MyoII in maintaining the spherical organization of the niche. Furthermore, the similarity in phenotype between ROK and MyoII inhibition suggests an essential role for AMC in niche shape.

**Figure 3:**
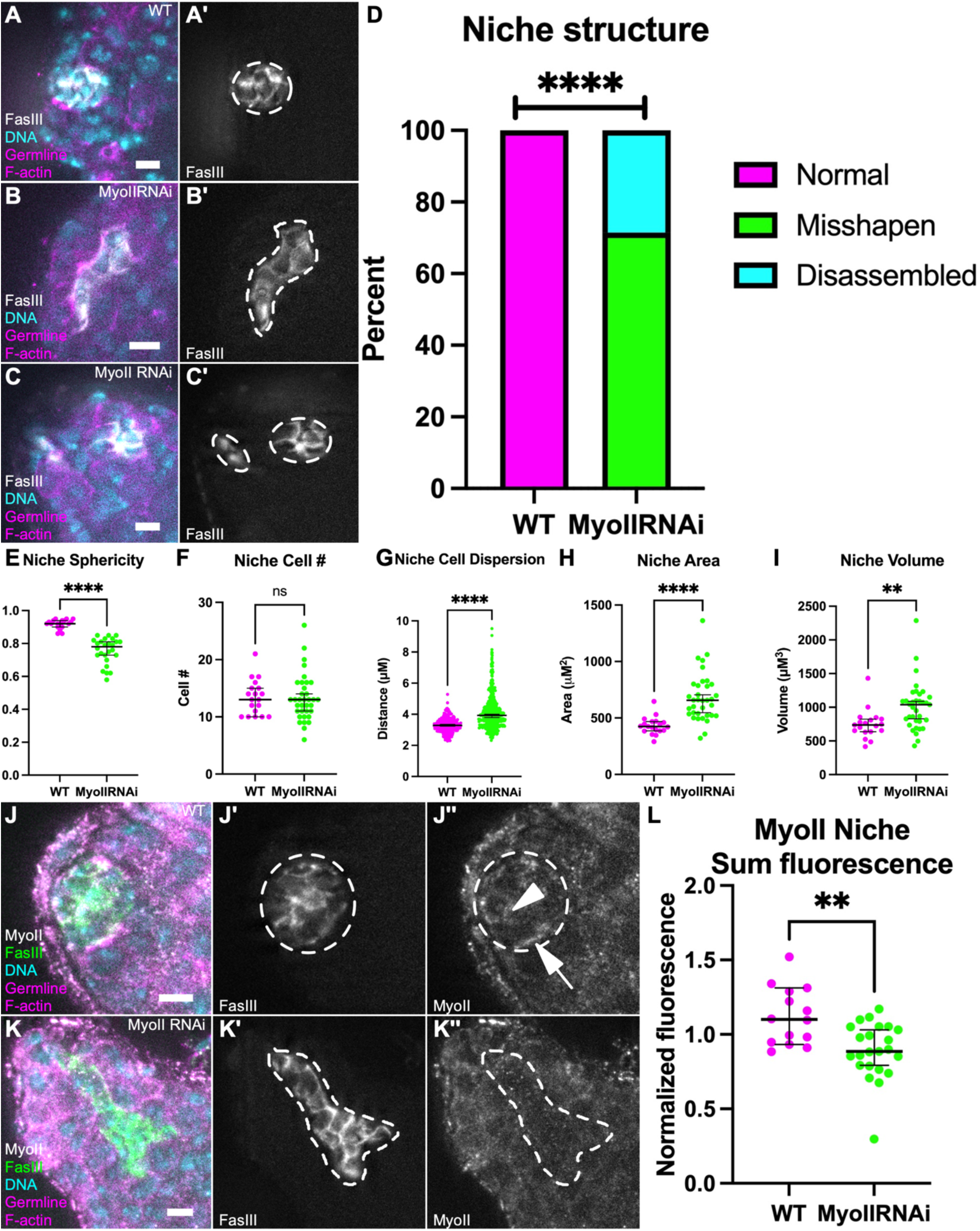
MyoII is required for niche shape. (A-C) Immunostained WT testis (A) and MyoIIRNAi testis where niche is misshapen (B) or disassembled(C); (A,B,C) nanos-lifeactdTom, FasIII (white), Hoescht (cyan); (A’,B’,C’) FasIII only; niche shape represented by white dotted line. (D) Quantification of niche disassembly shows MyoII RNAi niches are severely misshapen and, in some cases, disassembled into more than one niche aggregate compared to WT niches (WT: 0/19 niche misshapen or disassembled, MyoIIRNAi: 25/35 niches misshapen, 10/35 niches disassembled, ****p<0.0001, Chi-squared test). (E) Quantification of niche sphericity between WT and MyoIIRNAi niches (****p<0.0001, Mann-Whitney test). (F) Quantification of niche cell number show no difference between WT and MyoIIRNAi niches (ns). (G) Quantification of the average distance of a niche cell and its 3 nearest neighbors show increased distance between MyoIIRNAi niche cells compared to WT (****p<0.0001, Mann-Whitney test). (H-I) Quantification of the niche area (H) and volume (I) shows an increase in MyoIIRNAi niches compared to WT (****p<0.0001, **p=0.0010, Mann-Whitney test). (J-K)) Immunostained WT testis (J) and MyoIIRNAi testis (K); (J,K) nanos-lifeactdTom, FasIII (green), MyoIIHC (white); (J’,K’) FasIII only; (J’’,K’’) MyoIIHC only, MyoII localizes to niche-niche interfaces (white arrowhead, J’’) and also to niche-GSC interfaces (white arrow, J’’) in WT testes and this localization if abolished in MyoIIRNAi testes (K’’); niche shape represented by white dotted line. (L) Quantification of the sum of MyoIIHC fluorescence in the niche in MyoIIRNAi niches compared to WT niches (**p=0.0016, Mann-Whitney test).

### Niche structure is progressively disrupted upon MyoII inhibition

The strikingly separated niche fragments upon MyoII challenge suggested the possibility that, over time, constituent niche cells were drifting apart. To test this, we examined niches in which MyoII was compromised for a shorter period. These niches were still misshapen, though not as severely as in 10-day knockdowns including no disassembled niche fragments (Fig 4A-C). This was confirmed by sphericity measurements, where the 3-day knockdowns were significantly less spherical than wildtype, but not as misshapen as with 10 days of MyoII inhibition (Fig 4D). This suggests a progressive loss of niche structure the longer MyoII is compromised.

**Figure 4:**
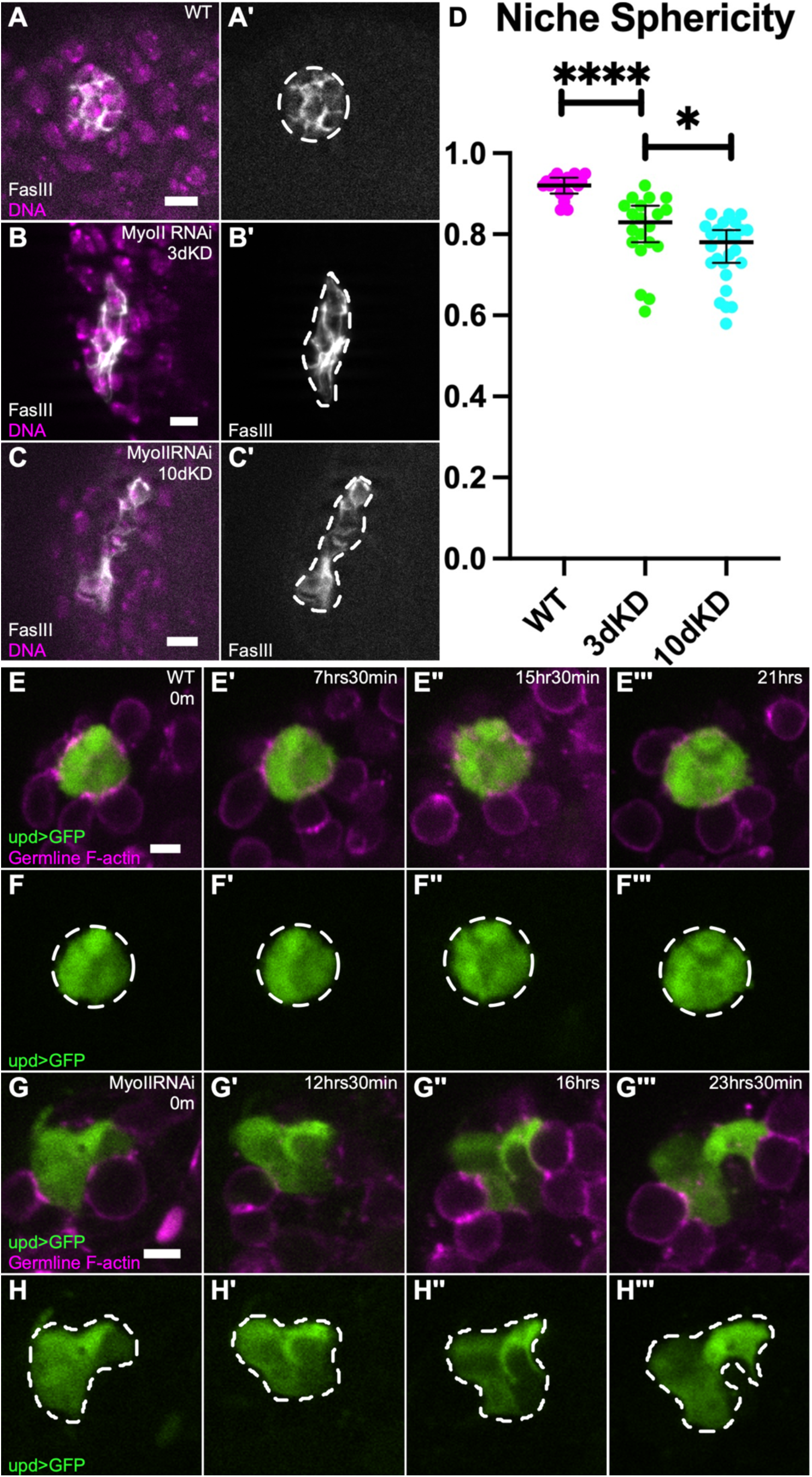
Niches progressively lose sphericity upon MyoII depletion. (A-C) Immunostained WT testis (A), and MyoIIRNAi testis, 3-day knockdown (B) and 10-day knockdown (C); (A,B,C) FasIII (white), Hoescht (magenta); (A’,B’,C’) FasIII only; niche shape represented by white dotted line. (D) Quantification of niche sphericity between WT and MyoIIRNAi 3 day and MyoIIRNAI 3 day and 10 day knockdown niches (****p<0.0001, *p=0.0157, Mann-Whitney test). (E-H) Time-lapse of WT niche (E,F) and MyoIIRNAi niche (G,H); (E,G) upd>GFP (green), nanos-lifeacttdTom (magenta); (F,H) upd>GFP only, dotted line represents niche shape at each time point.

To determine the mechanism by which MyoII inhibition disrupts niche structure, we performed live imaging to directly visualize niche cell behavior. Testes expressing MyoII RNAi in niche cells for 10 days were imaged *ex vivo* for a 24-hour period. We found that control niches generally maintained their compact shape throughout the imaging period, with the niche cells in the z plane shown remaining in a similar plane throughout imaging. Although membranes of individual niche cells were not always differentiated, there were no obvious changes in cell contacts to other niche cells or GSCs throughout imaging (Fig 4E-F). In contrast, niches compromised for MyoII progressively changed shape throughout imaging beyond their initial misshaped appearance. For example, niche cells in the MyoII knockdown seem to extend outward, potentially exchanging a previous contact with a neighbor niche cell, as well as making a new contact by surrounding a germline cell (Fig 4G-H). Additionally, niche cells can dynamically extend and retract protrusions in between germ cells (Fig S4, yellow arrows, white arrowheads). Together, this shows a progressive phenotype where niche cells compromised for MyoII are more loosely organized, and even explore surfaces of other cells.

### MyoII inhibition affected the polarized distribution of adhesion proteins between niche cells

In niches compromised for MyoII activity, the decrease in sphericity, increase in cell dispersion, and the behaviors visible by live-imaging all suggest that individual niche cells in the collective are not as tightly adherent, and can separate from one another. Elegant serial EM reconstruction showed that cells of the niche are extensively interdigitated with each other (Hardy et al., 1979). Subsequent analyses have revealed enrichment of proteins at niche-niche interfaces falling into several different adhesion classes, including E-Cadherin and N-Cadherin (Brower et al., 1981; Le Bras and Van Doren, 2006; Michel et al., 2011). We therefore asked whether loss of MyoII might be compromising the enrichment or polarity of these factors. In control testes, both E-cadherin and N-Cadherin exhibited enrichment at the niche as expected (Fig 5 A”,C”). Both proteins significantly polarized to the niche-niche compared to niche-GSC interfaces(Fig 5E,G). In contrast, when MyoII was compromised, the polarity of E-Cadherin and N-cadherin to niche-niche interfaces was significantly reduced, thought to a lesser degree for N-cadherin (Fig 5F,H).

**Figure 5:**
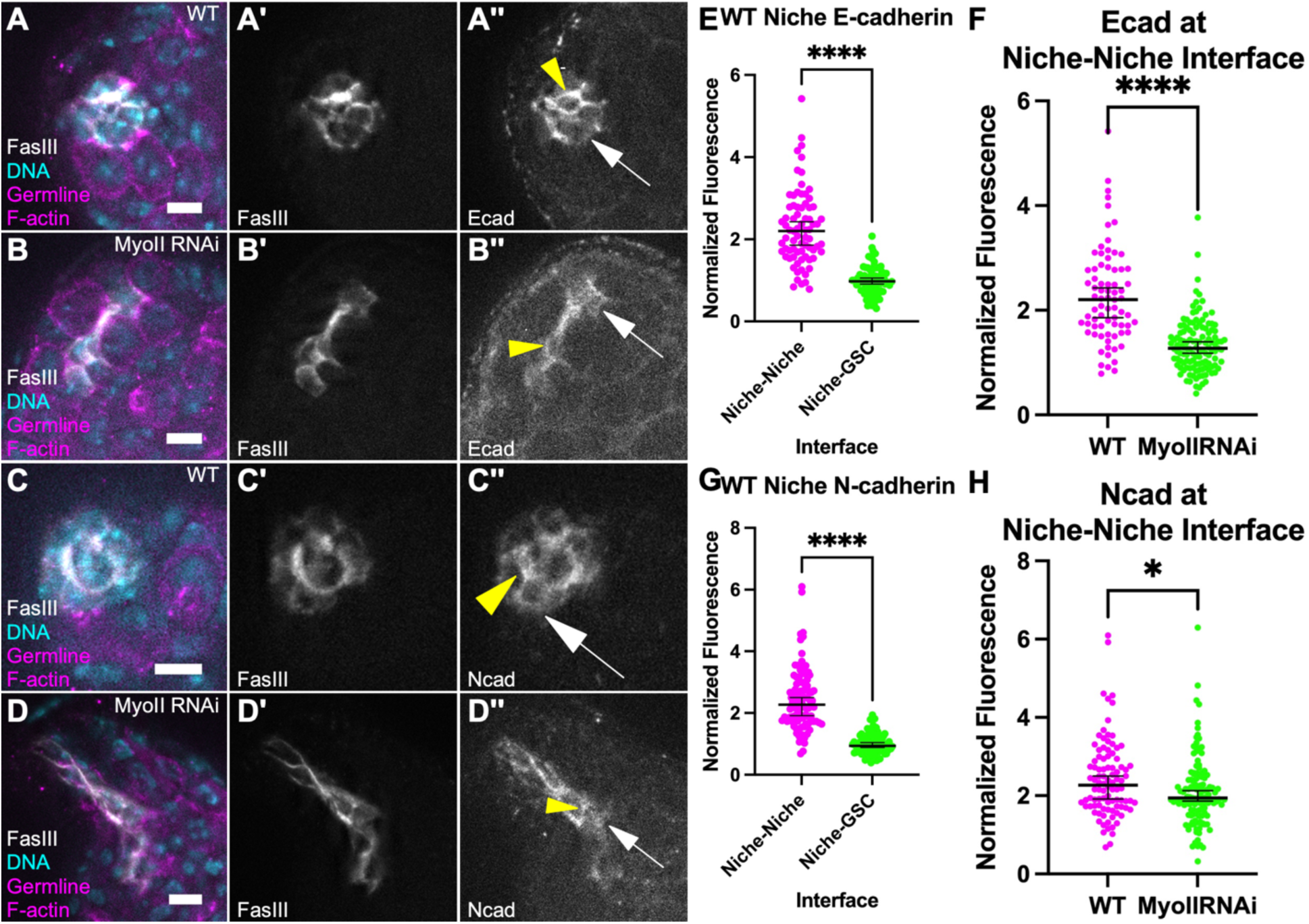
Disruption to polarized adhesion complexes when compromised for MyoII. (A-B) Immunostained WT testis (A) and MyoIIRNAi testis (B); (A,B) nanos-lifeacttdTom, FasIII (white), Hoescht (cyan); (A’,B’) FasIII only; (A’’,B’’) E-cadherin only; yellow arrowhead points to a niche-niche interface, white arrow point to niche-GSC interface. (C-D) Immunostained WT testis (C), and MyoIIRNAi testis (D); (C,D) nanos-lifeacttdTom, FasIII (white), Hoescht (cyan); (C’,D’) FasIII only; (C’’,D’’) N-cadherin only; yellow arrowhead points to a niche-niche interface, white arrow point to niche-GSC interface. (E,G) Quantification of WT E-cadherin (E) and N-cadherin (G) enrichment at Niche-Niche interfaces (A’’,C’’, yellow arrowheads) in comparison to Niche-GSC interfaces (A’’,C’’, white arrows) (****p<0.0001, Mann Whitney test). (F,H) Quantification of E-cadherin (F) and N-cadherin (H) enrichment at Niche-Niche interfaces (yellow arrowheads) normalized to Niche-GSC interfaces (white arrows) in MyoIIRNAi niches (B’’,D’’) compared to WT (A’’,C’’) (****p<0.0001,*p=0.0118, Mann Whitney test).

To test whether compromising E-cadherin was sufficient to disrupt niche structure, we expressed an RNAi against E-cadherin in the niche. Consistent with prior experiments, we observed no disruption in niche structure (Fig S5; (Michel et al., 2011)). As a more stringent test for the role of adherens junctions, we expressed an RNAi against beta-catenin(arm). Beta-catenin is likely integral to both E-cadherin and N-cadherin linkages and, thus, upon knockdown would compromise all adherens junctions (Oda et al., 1994). Similarly to compromising E-cadherin, we observed no disruption of the niche structure (Fig S5), even though beta-catenin was significantly reduced. The apparent stability of the niche when adherens junctions are compromised suggests that compromising MyoII affects not only cadherin complexes, but other adhesive complexes that must be acting redundantly among niche cells.

### Niches compromised for MyoII are functionally defective

The generation of misshapen and disassembled niches that still retain normal niche cell number allows us to ask how niche architecture might affect stem cell function. One of the key signals emanating from the testis niche, Unpaired (Upd), activates the Jak-STAT pathway only locally, in nearby somatic and germline cells. These first-tier cells represent the two stem cell lineages, CySC and GSC, and the restriction of signaling allows the progeny of these stem cells to differentiate properly. To determine whether niche signaling is compromised in misshapen niches, we immunostained for STAT in GSCs, as a readout for efficient Jak-STAT signaling. We observed that GSCs adherent to niches compromised for MyoII activate the Jak/STAT pathway; however they do so to a lesser degree than controls, suggesting inefficiency in signaling (Fig 6A-B,E). It is known that decreased STAT activation can cause GSC loss from the niche (Brawley and Matunis, 2004; Kiger et al., 2001; Leatherman and DiNardo, 2010; Tulina and Matunis, 2001). We therefore used live-imaging to directly track GSC behavior in control and MyoII-compromised niches. We scored loss events as germline cells that were initially adherent to niche but that subsequently lost contact over the imaging period (see Materials and Methods). We found that there was an increase in GSC loss from niches compromised for MyoII compared to controls (Fig 6C-D,G). Together these data show that misshapen niches have reduced Jak/STAT signaling and increased GSC loss.

**Figure 6:**
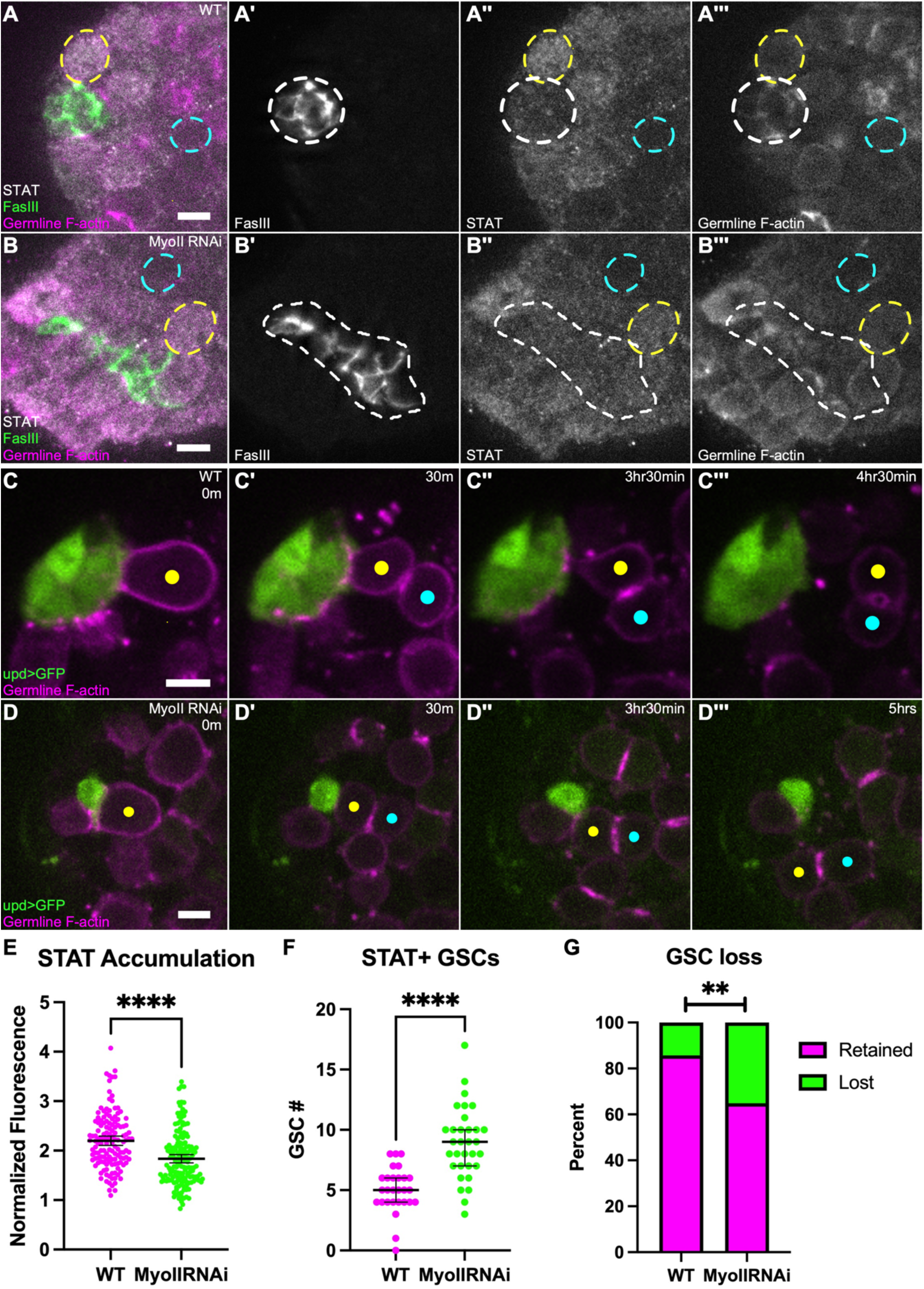
Niches compromised for MyoII are functionally defective. (A-B) Immunostained WT testis (A) and MyoIIRNAi testis (B); (A,B) nanos-lifeactdTom, FasIII (green), STAT (white); (A’,B’) FasIII only; (A’’,B’’) STAT only; (A’’’,B’’’) nanos-lifeacttdTom only; dotted white line indicates niche shape, dotted yellow circle indicates GSC, dotted blue circle indicates germ cell. (C-D) Time-lapse of relatively rare WT GSC loss (C) and more frequent MyoIIRNAi GSC loss (D), upd>GFP (green), nanos-lifeacttdTom (magenta), yellow dot indicates a GSC, blue dot indicates newly formed gonialblast; in both cases, GSC has oriented division (C’,D’), GSC-gonialblast (Gb) is eventually lost from the niche (C’’’,D’’’). (E) Quantification of STAT fluorescence intensity levels of GSCs in WT testes and MyoIIRNAi testes (****p<0.0001, Mann Whitney test). (F) Quantification of STAT+ GSCs in WT testes and MyoIIRNAi testes (****p<0.0001, Mann Whitney test). (G) Quantification of GSC loss in WT testes and MyoIIRNAi testes; WT: 10/70 GSCs, MyoIIRNAi: 40/114 GSCs (**p=0.0020, Fisher’s exact test).

The increase in area for niches compromised for MyoII (Fig 3H) raised the possibility that these disrupted niches might present more vacancies to attract cells to a stem cell fate. Therefore, we also scored the number of STAT-positive germ cells that contacted niche cells. We first chose a threshold signal intensity above which a cell would be scored as STAT-positive (see Materials and Methods). We found that there is a significant increase in the number of STAT-positive germ cells surrounding those niches compromised for MyoII than control niches (Fig 6F). These data together show that proper niche shape is important for restricting niche signals to a select population of cells, with disrupted niche structure increasing stem cell number but diminishing their ability to retain contact with the niche.

### More frequent symmetric outcomes of GSC divisions drive increased germ cells at MyoII knockdown niches

There are a few ways GSC number might increase at the niche. The first possibility is an increased GSC cell cycling rate. However, an EdU pulse ruled out any change in S phase frequency in GSCs between MyoII knockdowns and controls (Fig S2). The other possibilities each involved alterations in GSC divisions. One possibility was an increase in executing a ‘symmetric renewal’ event. Normally, GSCs divide perpendicular to the niche producing one daughter that renews as a stem cell and one gonialblast daughter moved farther from the niche that will eventually abscise and differentiate (Yamashita, 2003; Yamashita et al., 2007; Lenhart and DiNardo, 2015; Lenhart et al., 2019). A symmetric renewal can occur when, after a properly oriented division, the gonialblast daughter rotates inward and contacts the niche (Sheng and Matunis, 2011). The second way GSC number might increase is through a ‘symmetric division’, where the orientation is incorrect and the GSC divides parallel to the niche such that the two daughter cells remain in contact with the niche through and after division. We hypothesized that GSCs adherent to a misshapen niche might execute symmetric renewals and/or symmetric divisions at a higher frequency than in controls.

To test for symmetric renewal frequency, we live imaged GSCs in control testes and testes with depletion of MyoII from the niche and quantified the number of correctly oriented GSC divisions that were followed by contact of the gonialblast daughter with the niche. We observed an increase in symmetric renewal events at niches compromised for MyoII (Fig 7A-C,F). In these cases, the prospective gonialblast would appear to rotate in to contact the niche, although in some instances a niche cell appeared to extend outward to contact the daughter cell (Fig 7C’’’, white arrowhead). Therefore, niches compromised for MyoII allow for more cells to respond to niche signaling by an increase in symmetric renewal events. To test for symmetric divisions, we live imaged MyoII knockdown testes compared to controls and tracked each GSC division. While all GSCs in controls exhibited oriented divisions, about one quarter of GSC divisions in MyoII knockdowns were misoriented. These would result in both daughter cells adherent to the niche (Fig 7D-E,G). Taken together, niches compromised for MyoII have less efficient STAT activation and significant deviation from proper stem cell division control.

**Figure 7:**
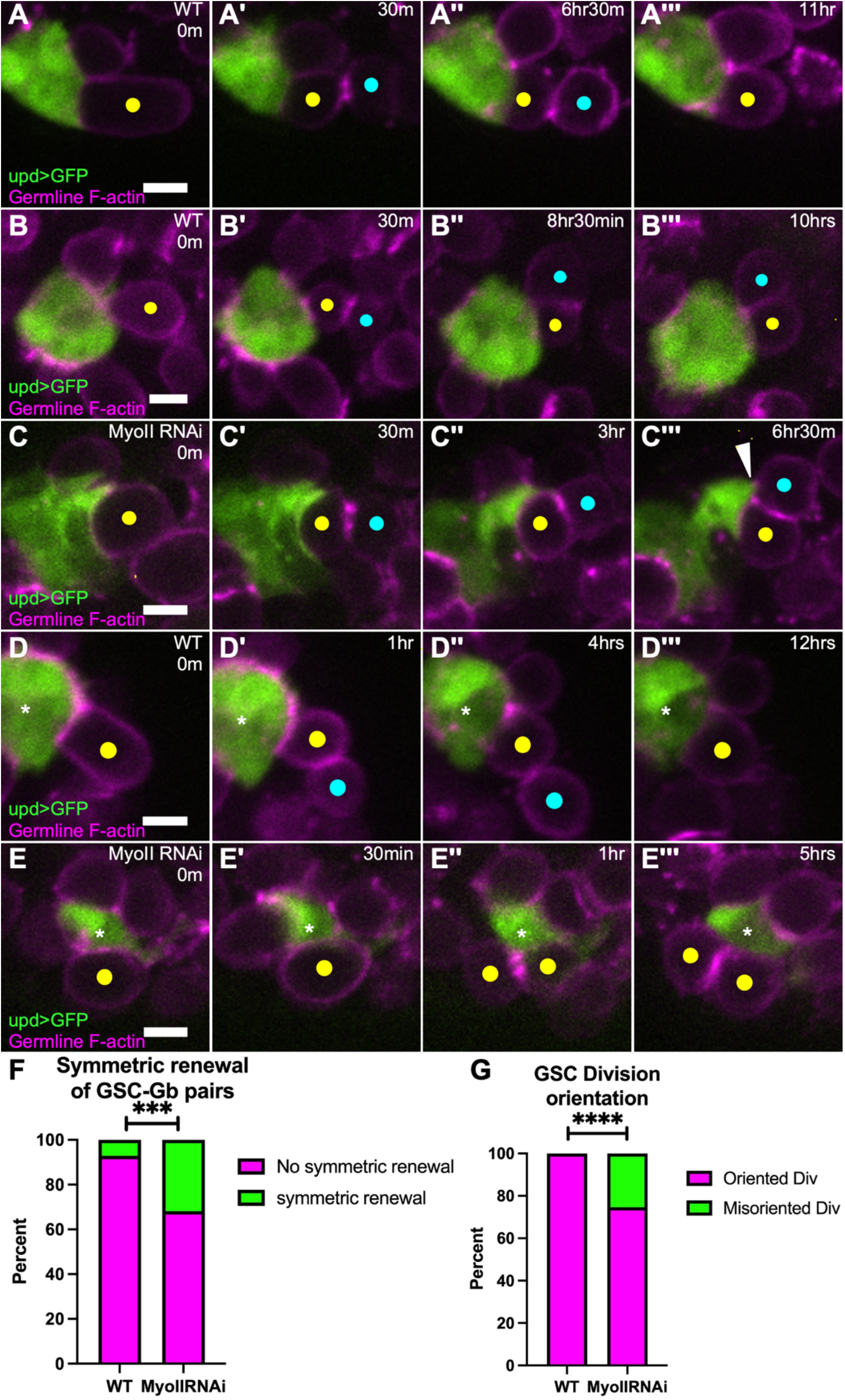
Germ cells repopulate the niche in MyoII knockdown niches. (A-C) Time-lapse of GSC divisions, upd>GFP (green), nanos-lifeacttdTom (magenta), yellow dot indicates a GSC, blue dot indicates newly formed gonialblast; (A) WT niche, GSC has oriented division; (B) WT niche, GSC has oriented division, but the newly generated GSC-Gb pair exhibits relatively rare symmetric renewal, and the gonialblast makes contact with the niche; (C) MyoII RNAi niche, GSC has oriented division, white arrowhead indicates niche extending outward to “capture” gonialblast for symmetric renewal (C’’’). (D-E) Time-lapse of WT testis (D) and MyoIIRNAi testis (E); upd>GFP (green, niche), nanos-lifeacttdTom (magenta, germ cells); in WT, GSC (yellow dot) divides perpendicular to niche (D’) and produces a GSC daughter (D’, yellow dot) and gb daughter (D’, blue dot) that will differentiate; in MyoIIRNAi, GSC (yellow dot) divides symmetrically to niche (E’) and produces a 2 GSC-like daughters, adherent to niche (E’, yellow dots). (F) Quantification of symmetric renewal events of GSC-Gb pairs in WT testes and MyoIIRNAi testes; WT: 6/82 GSC-Gb pairs, MyoIIRNAi: 29/93 GSC-Gb pairs exhibited symmetric renewal (****p<0.0001, Fisher’s exact test). (G) Quantification of division orientation of GSCs in WT and MyoIIRNAi testes; WT: 0/82 GSCs, MyoIIRNAi: 33/126 GSCs exhibited misoriented divisions (****p<0.0001, Fisher’s exact test).

### Niche MyoII is required for centrosome orientation checkpoint in GSCs

GSCs orient their divisions anchoring one centrosome at the niche GSC interface resulting in a spindle orientation that pushes the distal daughter cell away from the niche. A GSC-specific checkpoint prevents M phase entry if centrosomes are misoriented and that ‘centrosome orientation checkpoint’ (COC) is only released when a centrosome is re-captured properly (Cheng et al., 2008; Venkei and Yamashita, 2015). Because we observed symmetric divisions in MyoII knockdown niches, we hypothesized that this checkpoint was being bypassed in GSCs. We first tested whether there was centrosome misorientation using γ-tubulin as a centrosome marker. Indeed, we observed an increase in misorientation in GSCs with MyoII knockdown niches compared to aged-matched controls (Fig 8). A few GSCs appeared ‘double oriented’, as each centrosome was contacting a niche-GSC interface, suggesting that the subsequent division could result in two daughter cells adherent to a niche cell (Fig 8A-D).

**Figure 8:**
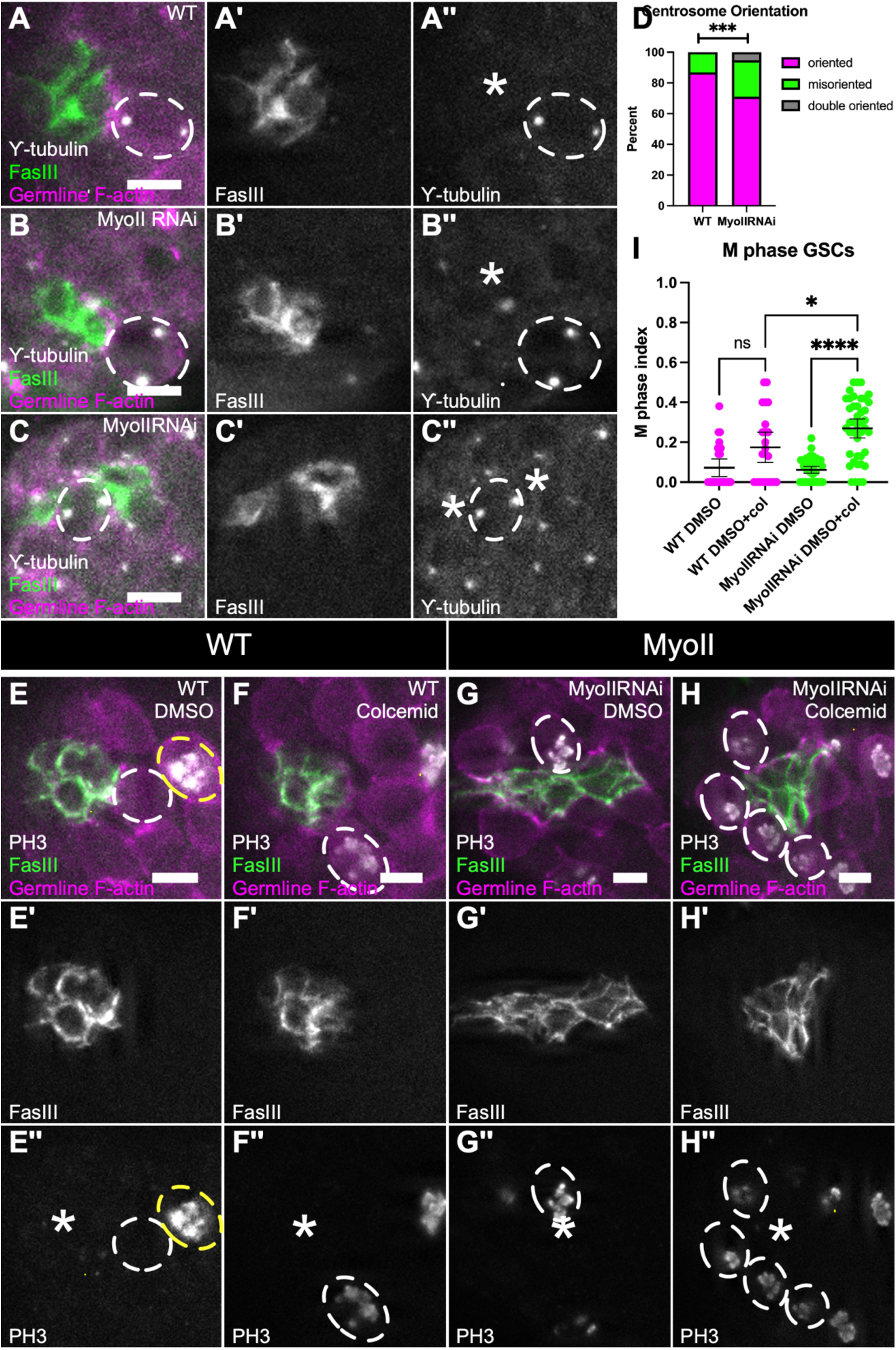
Niche MyoII required for centrosome orientation checkpoint in GSCs. (A-C) Immunostained WT (A) and MyoIIRNAi testes (B,C); (A,B,C) nanos-lifeactdTom, FasIII (green), γ-tubulin (white); (A’,B’,C’) FasIII only; (A’’,B’’,C’’) γ-tubulin only; white dotted lines indicate GSC, asterisk indicates niche. (D) Quantification of centrosome anchoring in GSCs in WT testes and MyoIIRNAi testes; WT: 15/114 GSCs with misanchored centrosomes, MyoIIRNAi: 113/476 GSCs with misanchored centrosomes, 25/476 GSCs with double anchored centrosomes (***p=0.0010, Chi-square test). (E-H’’) DMSO (E-E’’, G-G’’) and colcemid treated testes in WT (E-F’’’) and MyoII RNAi (G-H’’) conditions. (E,F,G,H) nanos-lifeactdTom, FasIII (green), PH3 (white); (E’,F’,G’,H’) FasIII only; (E’’,F’’,G’’,H’’) PH3 only; white dotted circle indicate GSC, asterisk indicates niche, yellow dotted circle indicates PH3 positive germ cell. (I) Quantification of M phase GSCs in WT and MyoII RNAi testes treated with DMSO or Colcemid; WT DMSO (n=26), WT DMSO+ colcemid (n=23), MyoIIRNAi DMSO (n= 45) MyoIIRNAi DMSO+ colcemid (n=42), (****p<0.0001,*p=0.0436, Kruskal-Wallis test).

The centrosome misorientation suggested that the COC might be defective in GSCs adherent to MyoII compromised niches. The established assay for COC activation takes advantage of differences between typical cells and GSCs in terms of when the cell cycle stalls following forced microtubule disassembly. Treatment with colcemid induces microtubule depolymerization which eliminates cytoplasmic asters necessary for centrosome anchoring, as well as mitotic spindles (Venkei and Yamashita, 2015). In most cycling cells, the absence of a bipolar spindle will cause arrest in M phase due to the spindle assembly checkpoint present in all cells (Musacchio and Salmon, 2007). This M-phase arrest is detected as an increased proportion of cells containing phosphorylated histone H3 (PH3). Under the same conditions, testis GSCs uniquely arrest *before* M phase due to loss of centrosome anchorage and activation of the COC, which occurs in G2 before the spindle assembly checkpoint (Venkei and Yamashita, 2015). Thus, loss of COC function can be identified as failure of GSCs to arrest in G2 and instead now arresting in M, indicated by an increased proportion of GSCs positive for PH3.

To assess the status of the COC, we treated control testes and testes with MyoII-compromised niches with colcemid and quantified the GSC M-phase index. In controls, we found no significant difference in M-phase index between vehicle- and colcemid treatments, as expected for GSCs with an active COC (Fig 8E-F,I). However, with MyoII-compromised niches there was a significant increase in M-phase positive GSCs after colcemid treatment (Fig 8G-H,I). This demonstrates that the COC is not functional in these GSCs. Taken together, these data suggest that MyoII-compromised disruptions to niche structure interfere with the establishment or activation of the COC in the adjacent germ cells.

## Discussion

We have shown that proper structure of the testis niche is required for its proper function. Disrupting the normally organized niche cell cluster leads to changes in signaling to stem cells, decreased retention of GSCs at the niche, and key changes in their tightly orchestrated division orientation program. Each of these alterations would be detrimental to robust spermatogenesis.

### ROK and MyoII are required for niche shape

AMC has been shown to induce cellular movements and migrations that can ultimately lead to specific organizations of cells. This is the case for the organization and/or placement of niches within tissues. In the formation of the intestinal crypt, contractility is required for its invagination (Sumigray et al., 2018). Our lab has recently shown that the early shaping of the male gonadal niche is dependent on AMC (Warder et al., 2023). Intriguingly, the continued enrichment of MyoII at the niche-stem cell interface in the adult testis coupled to the enrichment of ROK predicted by adult testis snRNA-seq data, suggested that contractility would play a key role in also maintaining niche architecture (Li et al., 2022; Raz et al., 2023).

We found that both ROK and MyoII are polarized at the niche-GSC interface. Reducing MyoII in niche cells led to decrease of MyoII signal suggesting that most of the MyoII contribution was by niche cells and not the germ cell cortex. The polarization of AMC components is initially established in the early embryo during compaction, where it generates tensile forces at the niche-GSC boundary (Warder et al., 2023). The ROK and MyoII enrichment at this boundary in the adult suggests that this tension remains at the niche-GSC interface to maintain the niche in a spherical organization. Indeed, we showed that knockdown of MyoII within niche cells severely disrupts niche structure. Importantly, loss of germ cells *does not* significantly alter niche architecture (Brawley and Matunis, 2004). Together, this suggests that preservation of AMC at the niche-GSC boundary from embryogenesis through adulthood is essential to maintain niche structure and that this polarization is primarily driven by the niche cells.

We show that absence of ROK and MyoII can lead to severe consequences to niche structure that ultimately cause defects in the neighboring GSCs. Since depletion of ROK and the MyoII heavy chain (zip) generated similar defects, we conclude that AMC is regulated by ROK in the cortex of cells, specifically along the niche-GSC interface. However, the lower frequency of niches with misshaping defects and the lesser degree to which niches are misshapen suggests other regulators also act in maintaining AMC at the niche. It is also possible that incomplete knockdown of ROK by RNAi might account for the difference in severity of the effect. In any case, our observations raise questions about how contractility factors are enriched in niche cells, how they are polarized to the correct interface with stem cells, and what the specific effects of cortical contractility might be that, when compromised, disrupt niche cell function, and generate substantial defects in stem cell behavior.

A potential source for candidate factors acting upstream to regulate AMC is the snRNAseq dataset of genes whose expression is enriched in niche cells. It is also possible that extrinsic signaling could help maintain niche shape. Interestingly, the initial architecture of this niche is shaped by feedback from recruited GSCs (Warder et al., 2023). Some of that feedback appears to be in the form of mechanical forces generated by GSC divisions (Warder et al., 2023). In the adult testis, the ablation of germ cells does not appear to alter niche shape (Brawley and Matunis, 2004), suggesting that, if extrinsic factors or forces exist, these cannot emanate from GSCs. Manipulations in the CySCs as well as manipulations in niche cells themselves, have been shown to affect niche architecture. However, the manipulations involved in those studies also caused changes in fate and proliferation of niche cells (Hétié et al., 2014; Greenspan and Matunis, 2018). Thus, it is not clear that CySCs specifically affect the cellular organization of the niche.

### Niche AMC maintains junctional complexes within the testis niche to promote adhesion between niche cells

We found that the niche progressively loses its shape when compromised for MyoII over time. Live-imaging first hinted at this, and afforded a unique perspective to understand niche shape changes, something typically challenging in other systems. Examining the effects of knockdown after different lengths of time revealed that the niche cells spread apart and suggested the possibility that adhesion was progressively compromised. Indeed, we observed changes to cadherin polarization when MyoII was compromised. Normally, the niche polarizes both E- and N-cadherin to the niche-niche interface as opposed to the niche-GSC interface. Interestingly, this polarization is opposite to that observed for AMC factors that are enriched at niche-GSC interfaces. The mechanism underlying this polarity opposition for these niche cells is not yet clear. However, this is a common phenomenon in other polarized tissues, such as epithelia (Simões et al., 2010; Bulgakova et al., 2013). There, a cascade involving microtubules, Par3 and RHO-GEF help maintain the enrichment of E-Cadherin to interfaces opposite to those with high contractility (Bulgakova et al., 2013). When these factors are compromised, epithelial organization is also compromised. Given that we observe niche cells drifting apart it will be of interest to test whether a similar cascade affects niche cell polarization.

It is likely that MyoII and contractility in niche cells affects more than cadherin polarization. Knocking down E-cadherin, or even beta-catenin/armadillo which likely challenges both E- and N-cadherin complexes, did not cause defects in niche shape (Michel et al., 2011). Indeed, a survey of the literature and the snRNA-sq data reveals a large number of junctional, adhesive and transmembrane factors that are enriched on these niche cells, such as Echinoid, Fasciclin III, Discs Large, and Polychaetoid (Raz et al., 2023). This suggests significant redundancy to the maintenance of niche organization. It will be of interest to pursue precisely which complexes are affected, and how, when AMC is compromised.

### Niche shape is required for efficient Jak/STAT signaling and proper GSC number

We observed that changes to niche structure had two effects on Jak/STAT signaling. First, we found that GSCs respond less robustly to Jak/STAT signaling when MyoII is compromised in the niche. It is known that GSCs unable to activate STAT properly are lost from the niche and we confirmed an increase in GSC loss when niche structure was disrupted (Kiger et al., 2001; Tulina and Matunis, 2001; Brawley and Matunis, 2004; Leatherman and DiNardo, 2010). Thus, the decreased GSC responsiveness to Jak/STAT signaling upon disruption of niche structure directly impacts a critical feature of GSC function (persistent adhesion to the niche).

Second, we found that there was an increase in the number of STAT-responding germ cells at MyoII-compromised. One hypothesis is that the area increase of the niche allows the signal to disperse more, which could afford more cells access to the signal, albeit each to a lesser degree. However, this cannot be confirmed without directly observing secreted ligand levels upon disruption of niche structure. Alternatively, there is a possibility that the activation of STAT itself is being disrupted upon loss of niche structure. There is evidence that STAT can be polarized to specific interfaces within the cell, and this polarization is required for proper STAT activation (Sotillos et al., 2008). Further studies could address whether STAT activation occurs at the interface in this system, and whether that is disrupted upon the misshaping or disassembly of the niche.

### Germ cells are retained at misshapen niches by executing symmetric behaviors

We found that more germ cells responded to STAT signaling when the niches were compromised for MyoII. It has been shown that there is competition for space at the niche, so it is not surprising that cells quickly fill a space that might be generated by increases to niche area and volume (Issigonis et al., 2009; Sheng et al., 2009). In principle, while de-differentiation of gonial cells could account for such an increase, this is a relatively rare event and was not obvious in our live-imaging. Instead, our imaging illuminated two behaviors in GSCs attached to compromised niches that could account for their increase in number at the niche. First, we observed GSCs that underwent a properly oriented division to push the gonialblast away from the niche. However, the still-attached gonialblast instead contacted the niche as opposed to remaining pushed away from the niche. This type of ‘symmetric renewal’ is normally a rare event but was significantly increased in MyoII-compromised niches (Figure 6J). In some cases, we observed niche cells that appeared to extend outward to contact the gonialblast cell (Figure 6E’’’). Whether this reflects some attraction or is purely a stochastic consequence of the aberrant niche shape remains to be investigated. Second, we observed a higher frequency of mis-oriented GSC divisions (discussed below). Such ‘symmetric divisions’ do not occur in GSCs normally. Thus, when niche organization is disrupted, both symmetric renewal and symmetric division occurs, leading to gonialblast daughter cells that are meant for differentiation, potentially maintaining a stem cell fate instead.

### Proper niche structure is required for proper stem cell division

In various tissues, stem cells divide in an oriented manner by precisely positioning of one or both centrosomes so that the proper orientation of the eventual spindle is established. For example, in *Drosophila* neuroblasts, centrosome asters are regulated such that one stabilizes at the apical cortex and the other travels to the opposite pole due to its loss of microtubule stability (Rusan and Peifer, 2007). In the case of the *Drosophila* testis, one centrosome in the GSC anchors at the niche-GSC interface to ensure a properly oriented division. Several factors have been identified that all act intrinsically in the GSC to anchor the centrosome. For example, it has been shown that there are polarity proteins like Bazooka/Par-3 and pericentriolar proteins like centrosomin that localize to the niche-GSC interface within the GSC and are required for proper centrosome orientation (Yamashita, 2003; Inaba et al., 2010; Inaba et al., 2015a). The oriented division is an important feature in this tissue, and as such GSCs uniquely engage a centrosome orientation checkpoint (COC), which will cause arrest in G2 if the centrosomes are not properly oriented for division (Cheng, 2008; Yuan, 2011; Venkei, 2015)(Cheng et al., 2008; Yuan et al., 2012b; Venkei and Yamashita, 2015). Significantly, in disrupted niches we observed mis-oriented centrosomes, and a failure of the checkpoint resulting in mis-oriented GSC divisions. This is the first description of a niche-dependent role in centrosome orientation and the subsequent execution of the COC in GSCs. We hypothesize that the cytoskeletal connections among MyoII, actin and adherens (or other junctional proteins) in the niche cell cortex must be what link to junctional factors within the GSC cortex to ensure proper centrosome orientation.

Taken together, this work emphasizes how AMC dependent niche structure is critical for proper stem cell behavior and ultimately necessary for maintaining the proper balance of stem cells within a tissue.

### Limitations of this study

Although our work shows preservation of niche structure through AMC, we were unable to distinguish the precise mechanism by which niche AMC maintains this structure. Additionally, our work shows a decrease in STAT response in GSCs with misshapen niches, however we could not directly assay ligand levels or distribution. Lastly, we determined that the COC is bypassed occasionally in GSCs with misshapen niches, however we could not precisely determine the mechanism by which the COC is inactivated.

## Materials and Methods

### RNAi experiments

Virgin females with updGal4 tubGal80ts and a UAS-GFP construct were crossed with males of the RNAi for that particular experiment. These crosses were kept at 18 degrees until late pupation, and were then upshifted to 29 degrees to induce expression of the RNAi and GFP. Males were then dissected after 10 days at 29 degrees unless otherwise indicated. For MyoII HC RNAi, E-cadherin RNAi, and Armadillo RNAi experiments, controls were aged matched and dissected at the same time as knockdown, and did not include the Gal4 driver. For ROK RNAi experiments, controls were kept at 18 degrees for 10 days after late pupation and had the same genotype as ROK knockdowns.

### Testis dissections and immunostaining

Testes were dissected in ringers and fixed in 4% formaldehyde in Buffer B (16.7mM KPO_4_, pH 6.8; 75mM KCl; 25mM NaCl; 3.3mM MgCl_2_), and 0.1% Triton for 20 minutes. They were then washed in PBS plus Triton for 20 minutes and then blocked in 4% normal donkey serum for one hour. Primary antibodies were diluted in block and added overnight at 4° C. Secondary antibodies were diluted in block and used at 3-4ug/ml for 1 hour at room temperature (Alexa488, Cy3, orAlexa647; Molecular Probes and Jackson ImmunoResearch). Hoechst at 0.2 ug/ml was used to stain DNA for 3 minutes. Testes were then equilibrated in 50/50 Ringers solution (5mM HEPES, pH 7.3; 130mM NaCl; 5mM KCl; 2mM MgCl_2_; 2mM CaCl_2_) and glycerol and mounted in 2% n-propyl-gallate, 80% glycerol.

We used chick antibody against GFP 1:3000 (Aves labs, GFP-1020), rabbit antibody against RFP 1:500 (Abcam, ab62341), Vasa 1:5000 (gift from R. Lehmann, MIT), STAT92E 1:375 (gift from E. Bach, NYU), MyoII HC (zipper) 1:750 (gift from D. Kiehart), Phosphohistone H3 (PH3) 1:1000 (Upstate Biotech, 06-570); mouse antibody against Fasciclin III 1:50 (DSHB, 7G10), Gamma Tubulin 1:200 (Sigma, GTU-88), Beta-catenin (armadillo) 1:100 (DSHB, N2 7A1), rat antibody against DE-Cadherin 1:20 (DSHB, DCAD2), DN-Cadherin (DSHB, Ex#8). Images of fixed testes were acquired on a Zeiss Imager with an apotome using a 40x, 1.2 N.A. water immersion lens.

### 3D Analysis

To quantify niche shape defects, we stained with Fasciclin III and evaluated in 3D whether each niche was normal, misshapen or disassembled using Imaris software. Normal niches were scored as such if the Fasciclin III staining resembled a compact, relatively spherical shape. Misshapen niches were scored as such if they were intact niches that had lost shape or were less compact, typically adopting elongate shapes. Disassembled niches were scored as such if they were fragmented. This type of categorization was used for MyoII knockdown and ROK knockdown experiments, with Chi-squared tests used to evaluate comparisons.

Imaris software was also used to extract sphericity, area, and volume of the niche. To extract these values we used Fasciclin III labeling of the niche to draw a surface that encompasses the whole niche. The surface was then generated into a mask and further refined by drawing a more accurate surface on the new mask that encompasses only the niche. This second surface was used to extract sphericity, area, and volume calculations. For all sphericity measurements, Fasciclin III was used except for when E-cadherin was used to draw surfaces for WT in Figure 1. Data was imported into Prism v10.0 to generate scatterplots where each dot represents a testis. To quantify niche cell number and average nuclear distance from one niche cell to its 3 nearest neighbors, we used Hoechst to label niche cell nuclei that were only within the confines of the finalized niche surface. This allowed us to accurately quantify niche cell number and the average distance between them. Data was imported into Prism v10.0 to generate scatterplots where each dot represents a niche cell nuclei. Mann-Whitney tests were used to evaluate comparisons.

### Live Imaging

Live Imaging was carried out as in Lenhart et al 2015. Control and MyoII RNAi samples were dissected after 10 days at 29 degrees and mounted on a matek dish. Imaging media was carefully added, which consisted of 15% FBS (Gibco 10082) and 0.5X penicillin/streptomycin (Corning 30-002-Cl) in Schneider’s Insect Media (Gibco 21720-024) (Prasad et al., 2007; Sheng and Matunis, 2011; Lenhart and DiNardo, 2015). The stock imaging media was prepared ahead of time, kept at 4 °C, and used within two weeks. Bovine insulin (Sigma l0516) was added to an aliquot of imaging media, just prior to use, to final concentration of 0.2 mg/mL. Upd>GFP labeled the niche and noslifeacttdTomato (Lin et al., 2020) labeled germ cells to assess niche changes and germ cell behaviors, respectively, throughout imaging. Live-imaging was carried out overnight using 30 minute intervals, and z step sizes of 0.5uM; stack depth was 40uM to include the whole niche. Testes were imaged on an Olympus iX83 with a Yokagawa CSU-10 spinning disk scan head, 60X 1.4NA silicon oil immersion objective, and Hamamatsu EM-CCD camera.

### Quantification of ROK, MyoII and Cadherin fluorescence

To quantify polarity of either ROK and MyoII between niche-niche interfaces and niche-GSC interfaces we examined transgenic lines for ROK (ROK-GFP), MyoII heavy chain (zip::GFP) and MyoII regulatory light chain (sqh3xGFP). We identified niche-niche interfaces by FasIII staining and identified niche-GSC interfaces by comparing FasIII staining to germ cell markers, which consisted of either a vasa stain or using the noslifeacttdTomato transgenic line. To characterize the polarity of either ROK or MyoII at the niche-GSC interface we used Image J to draw a 5 pixel-wide line along 3 niche-GSC interfaces and extracted the fluorescence intensity. These values were normalized to the average fluorescence intensity along 3 niche-niche interfaces, after background subtraction.

To quantify polarity of either E-cadherin and N-cadherin between niche-niche interfaces and niche-GSC interfaces we analyzed testes stained for either E-cadherin or N-cadherin, and Fasciclin III. We identified niche-niche interfaces by FasIII staining and identified niche-GSC interfaces by comparing FasIII staining to the germ cell marker, noslifeacttdTomato transgenic line. Using Image J, we measured the fluorescence intensity along 3 niche-niche interfaces and normalized them to the average fluorescence intensity along 3 niche-GSC interfaces, after background subtraction, to quantify the polarity of E-cadherin and N-cadherin at the niche-niche interface. We repeated these measurements in MyoII knockdown niches to determine whether there was a change in polarity of E-cadherin or N-cadherin in niche cells. Data was imported into Prism v10.0 to generate scatter plots with each dot representing an interface. Mann-Whitney tests were used to evaluate all comparisons.

### Quantification of MyoII knockdown

To quantify the level of MyoI knockdown, Myosin II knockdown and control testes were stained with an antibody against MyoII HC. Using Image J, we chose representative z-slice and drew an ROI that encompassed the niche, identified by FasIII staining. This same ROI was moved further along the testis sample, to a position among the germ cells for normalization. After background subtracting, the sum of fluorescence within the niche ROI was normalized to the sum of fluorescence of the germ cell ROI. This was done for each testis, and those values were imported into Prism v10.0 and scatterplotted for MyoII knockdowns and controls where each dot represents a testis. A Mann-Whitney test was used to evaluate the comparison.

### Quantification of STAT accumulation

To quantify the level of STAT accumulation, MyoII knockdown and control testes were stained with an antibody against STAT. Using Image J, ROIs were taken from all GSCs to extract mean fluorescence intensity from each cell. A cell was scored as a GSC if it was contacting the niche, as determined by using noslifeacttdTom for the germ cells and FasIII for the niche. Within each testis, 3 additional germ cells (non GSCs) were measured and averaged together after background subtraction to yield a ‘germ cell mean intensity’ value. Each GSC measurement within that testis was normalized to that germ cell mean fluorescence intensity. Data was imported into Prism v10.0 to generate scatterplots where each dot represents a GSC. A Mann-Whitney test was used to evaluate the comparison.

### Quantification of STAT positive GSCs

To determine whether a germ cell adherent to the niche was STAT positive, we quantified the mean fluorescence intensity from each cell that was adherent to the niche as described above. We determined the mean STAT enrichment for all cells measured for both the controls and MyoII knockdowns. We established individual thresholds for the control group as well as the MyoII knockdown group. We determined the threshold to be one standard deviation below the relative mean fluorescence intensity of controls. For controls, the relative mean fluorescence intensity was 2.18, with a standard deviation of 0.53, therefore any cells with a mean fluorescence intensity above 1.65 was counted as STAT positive. For MyoII knockdowns the relative mean fluorescence intensity was 1.89, with a standard deviation of 0.58, therefore any cell with a mean fluorescence intensity above 1.31 was counted as STAT positive. Data was imported into Prism v10.0 to generate scatterplots where each dot represents a testis. A Mann-Whitney test was used to evaluate the comparison.

### Quantification of Live Imaging Behavior

To analyze GSC behavior, each GSC was tracked from the start of imaging (t 0) until the end of the 24 hour imaging session. GSCs were scored for division orientation, symmetric renewal, and loss. GSCs that divided and retained one daughter cell at the niche and the other pushed away from the niche were scored as properly oriented. GSCs that divided such that both daughter cells remained in contact with the niche for multiple time points after division were scored as misoriented. The frequency of misoriented divisions relative to all divisions was plotted comparing MyoII knockdown niches and controls. A Fisher’s exact test was used to evaluate the comparison.

GSCs that had oriented divisions were simultaneously tracked for whether the GSC-Gonialblast (Gb) pair exhibited a symmetric renewal event. The criteria for such would mean that a newly produced Gb initially not touching a niche cell would ultimately rotate in and make contact with the niche and remain in contact for multiple time points. The frequency of symmetric renewal events out of every oriented division scored was plotted between MyoII knockdown niches and controls. A Fisher’s exact test was used to evaluate the comparison.

GSCs that were lost from the niche were scored as those cells initially touching a niche cell but ultimately losing contact and drifting away after multiple time points. The frequency of GSC loss out of all GSCs scored was plotted between MyoII knockdown niches and controls. A Fisher’s exact test was used to evaluate the comparison.

### Quantification of centrosome orientation

To determine whether centrosomes were properly oriented in MyoII knockdowns compared to WT testes, we stained for centrosomes using an antibody against γ-tubulin. We used Fasciclin III staining to visualize the niche and used noslifeacttdTom to visualize germ cells. GSCs were scored as having oriented centrosomes if they had 2 visible centrosomes with one touching the niche-GSC interface within the GSC. GSCs were scored as having misoriented centrosomes if they had 2 visible centrosomes, neither of which were touching the niche-GSC interface. GSCs were scored as having doubly oriented centrosomes if they had 2 visible centrosomes where both were touching a niche-GSC interface. A chi-squared test was used to evaluate the comparison.

### Quantification of S-phase in niche cells and GSCs

To quantify frequency of S-phase entry in GSCs and in niche cells, EdU pulses were done using the Click-iT EdU plus kit (Thermofisher, C10419) (Salic and Mitchison, 2008). Immediately after dissection in Ringers, testes were incubated in 10uM of EdU in Ringers for 30 minutes at room temperature before fixing. The azide reaction to couple EdU to Alexa 647 was carried out after fixing, but prior to the remainder of the staining protocol. GSCs positive for EdU were scored and the frequency of EdU positive GSCs out of all GSCs was plotted per testis between MyoII knockdown niches and controls. Data was imported into Prism v10.0 to generate scatterplots where each dot represents a testis. A Mann-Whitney test was used to evaluate the comparison. Niches were also screened for any EdU positive niche cells, but they were not observed.

### Colcemid Experiments

To determine whether the centrosome orientation checkpoint (COC) was active, we assayed for its activity using the protocol of Venkei and Yamashita, 2015. Immediately after dissection in Ringers, testes were transferred into imaging media plus insulin, containing either DMSO only as a control, or DMSO + 10mM colcemid, for a final concentration of 100μM. The samples were incubated in 4.5 hours at room temperature. Samples were fixed as usual and stained with a rabbit antibody against PH3 to score for cells in mitosis. GSCs positive for PH3 were scored and the frequency of PH3-positive GSCs out of all GSCs was plotted for MyoII knockdown niches and controls between colcemid treated and DMSO only conditions. Data was imported into Prism v10.0 to generate scatterplots where each dot represents a testis. A Kruskal-Wallis test with Dunn comparisons was used to evaluate all comparisons.

## Acknowledgments

We thank the Bloomington Drosophila Stock Center (NIH P40OD018537), Vienna Drosophila Research Center, R. Lehmann, E. Bach, Y. Bellaïche, D. Kiehart, and Zallen for antibodies and stocks. We thank the director of our CDB core for use of Imaris workstations. Thank you to V. Desikan on initial experimental set up. Thanks to B. Warder, K. Nelson and the Lenhart Lab for input throughout this project. Special thanks to Kari Lenhart for her manuscript comments and her microscope for which our live imaging experiments would not be possible.

## Competing interests

The authors declare no competing interests.

## Funding

GM008216 to G.S.V., GM136270 to S.D.

## Data Availability

All relevant data can be found within the article and its supplementary information.

## Notes

### Competing Interest Statement

The authors have declared no competing interest.

